# MskAge - An Epigenetic Biomarker of Musculoskeletal Age Derived from a Genetic Algorithm Islands Model

**DOI:** 10.1101/2023.05.04.539347

**Authors:** Daniel Charles Green, Louise Reynard, James Henstock, Sjur Reppe, Kaare Gutvik, Mandy Peffers, Daryl Shanley, Peter Clegg, Elizabeth Canty-Laird

## Abstract

**Background:** Age is a significant risk factor for functional decline and disease of the musculoskeletal system, yet few biomarkers exist to facilitate ageing research in musculoskeletal tissues. Multivariate models based on DNA methylation, termed epigenetic clocks, have shown promise as markers of biological age. However, the accuracy of existing epigenetic clocks in musculoskeletal tissues are no more, and often less accurate than a randomly sampled baseline model.

**Results:** We developed a highly accurate epigenetic clock, MskAge, that is specific to tissues and cells of the musculoskeletal system. MskAge was built using a penalised genetic algorithm islands model that addresses multi-tissue clock bias. The final model was trained on the transformed principal components of CpGs selected by the genetic algorithm, which are significantly enriched for pathways terms related to the skeletal system and mesenchyme development. We show that MskAge tracks epigenetic ageing *ex vivo* and *in vitro*. Epigenetic age estimates are rejuvenated to zero with cellular reprogramming and are accelerated at a rate of 0.45 years per population doubling. Remarkably, MskAge explains more variance associated with *in vitro* ageing of fibroblasts than the purpose-developed skin and blood clock.

**Conclusion:** The precision of MskAge and its ability to capture perturbations in biological ageing make it a promising research tool for musculoskeletal and ageing biologists.

## Introduction

Ageing is a complex and multifaceted process manifesting in a progressive decline of cellular function that is historically linked to the passing of time (1). While societal norms associate an individual’s age with the number of years since birth, the rate of organismal development and decline is non-linear across a lifespan and varies between tissues and organisms. This non-linearity and heterogeneity differentiate biological ageing trajectories from those typically associated with chronological ageing. Owing to the complexity of the ageing process, accurate biomarkers of biological age are becoming increasingly valuable in research and clinical settings (2, 3). Most existing biomarkers primarily focus on accessible tissues, such as blood, skin and saliva (2, 4, 5). Although these biomarkers are useful for investigating ageing rates at the population level, they may not be as applicable to tissues obtained through invasive procedures, such as those in the musculoskeletal system. To maximise the likelihood of successfully increasing health span and lifespan, interventions will likely need to rejuvenate cells throughout the entire organism. Consequently, identifying biomarkers that facilitate *ex vivo* research on musculoskeletal tissues has the potential to elucidate anti-ageing interventions currently beyond the scope of existing biomarkers.

In the past decade, DNA methylation (DNAm) clocks have emerged as some of the most promising and accurate biomarkers of age (2, 5–8). DNAm clocks are multivariate statistical or machine learning models that predict chronological age or age-related outcomes based on the proportion of methyl groups (CH_3_) present on DNA at an optimised selection of CpGs. Notably, CpGs selected by clocks are not necessarily those within ageing genes but rather those with associations to age in thousands of samples (7). The relative stability of DNA methylation allows DNAm clocks to achieve higher accuracy compared to biomarkers developed on other layers of molecular data (9). Moreover, the addition of a methyl group to cytosine by DNA methyl transferases (DNMTs) is reversible via its subsequent hydroxylation by the Ten Eleven Translocase (TET) enzymes, providing a dynamic mechanism that can be exploited to track and potentially modify the rate of biological ageing through the methylome (10, 11). The divergence between predicted epigenetic age and chronological age is referred to as epigenetic age acceleration (12). It has been well established that DNAm clocks can predict age acceleration rates independently of chronological age, age-related disease and age-related functional outcomes (e.g., hand grip strength), making them promising candidates as biomarkers of ageing (2, 4, 5, 7, 12–15).

Ageing profoundly impacts musculoskeletal function, leading to the deterioration of bone, cartilage and muscle, which can cause diseases such as osteoporosis and osteoarthritis, as well as the ageing syndrome of frailty (16, 17). Although attempts have been made to define biomarkers of ageing in musculoskeletal tissues, these primarily focus on biomarkers of bone turnover, skeletal muscle mass, or assessments of physical activity (18). While these biomarkers are useful for *in vivo* monitoring of physical function, they do not directly assess the age of musculoskeletal cells and cannot be applied *ex vivo*. To facilitate *ex vivo* research on musculoskeletal tissues, highly accurate and specific epigenetic biomarkers or other molecular clocks are likely required.

Some existing epigenetic clocks, such as Horvath’s original multi-tissue clock, are applicable to a wide range of tissue and cell types (7). However, the representation of musculoskeletal tissues in the training of these epigenetic clocks is limited compared to other tissues, making them less reliable for musculoskeletal ageing research due to the tissuespecific nature of DNA methylation. The primary drawback of non-specific epigenetic clocks is their poor accuracy and high variance in musculoskeletal tissues, which has implications for designing experiments with appropriate power. Obtaining musculoskeletal samples for research often involves highly invasive procedures, such as surgical operations or post-mortem extractions, highlighting the need for more accurate and specific epigenetic clocks to design appropriately powered studies. Recently, a highly accurate epigenetic clock (error +/- 4.6 years) was developed to track the age of human skeletal muscle using 19,000 CpGs common between the Infinium 27k, Infinium 450k and Infinium EPIC arrays (8). The skeletal muscle clock exemplifies an instance where the limitations of Horvath’s original multi-tissue clock were addressed, particularly its poor calibration for underrepresented samples in the training set of the model.

In the present study, we demonstrate that existing epigenetic clocks are no more and, in some instances, less accurate than a randomly sampled baseline model across multiple musculoskeletal tissues. We employed a genetic algorithm-based meta-heuristic feature selection framework to construct a musculoskeletal tissue-specific epigenetic clock, optimising CpG selection to minimise error within each tissue without bias from unequal tissue distributions common to multitissue clocks. Recently, it was shown that the algebraic transformation of CpG methylation into principal components for use as features in training an epigenetic clock addresses several technical issues arising from CpG feature models (19). Our final model, MskAge, uses a modified version of this framework, where principal component analysis is calculated after feature selection with the genetic algorithm. We demonstrate that, this modified approach, applied to our dataset, improves accuracy compared to calculating principal components of the entire feature space, resulting in a highly accurate epigenetic clock (error +/- 3.51 years) across all musculoskeletal tissues and blood samples in the dataset. Although MskAge is designed to predict chronological age, it resets across multiple independent cellular rejuvenation datasets and strongly correlates with population doubling *in vitro*. A tutorial for calculating MskAge can be found at Msk-Age Github.

## Results

### Acquired DNA Methylation Datasets

Infinium 450k and EPIC methylation array data were acquired from searches of the public domain repository Gene Expression Omnibus (GEO) and novel data generated within the UK Center for Integrated Musculoskeletal Ageing (CIMA). A total of 1,048 samples were identified across both platforms. Figure 1 shows an overview of the analysis pipeline for the generation of MskAge.

**Fig. 1.**
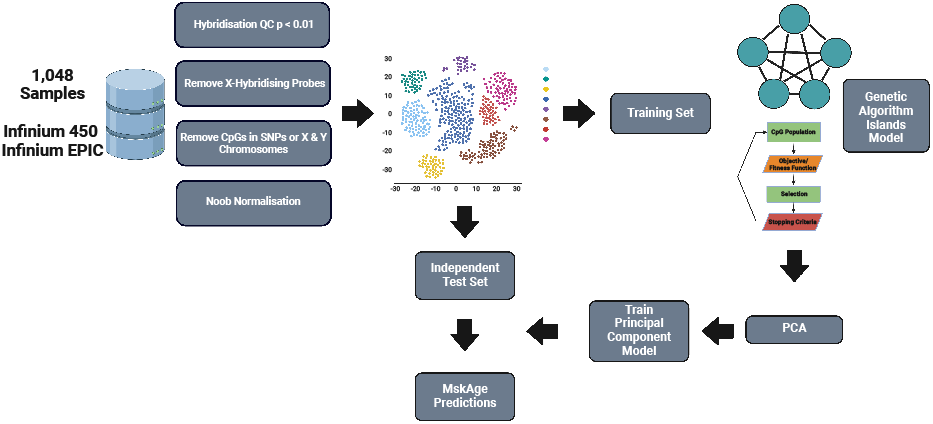
Workflow schematic. Schematic of analysis workflow for the development of Musculoskeletal Age (MskAge). The schematic depicts the collated samples undergoing quality control and filtering, being merged into a matrix and finally overviews the process of training the genetic algorithm and principal component model.

### Methods of Quantile Normalisation Skew Epigenetic Age Esitmates

With a plethora of normalisation methods available for methylation arrays technologies, the impact of preprocessing on epigenetic age estimates is an important consideration (20–23). To explore this issue and guide the choice of normalisation for MskAge development, we computed the epigenetic age of 16 cartilage samples with a relatively narrow age range (50 to 60 years) using different methods. Our analysis revealed that for PhenoAge, GrimAge, and Hannum clocks, the Quantile normalisation employed by both Quantile and Funnorm methods significantly affected epigenetic age estimates (Supplementary Figure 1). This effect is further highlighted by a correlation matrix computed for each respective normalisation method, demonstrating that Quantile normalisation methods produce distinct methylation profiles that cluster independently from non-quantile methods (Supplementary Figure 2). These methods lead to significantly exacerbated errors in epigenetic clocks not trained on quantile-normalised data (FDR *<* 0.001) (Supplementary Figure 1). Based on these results, we select single-sample normal exponential using out-of-band probes (ssNoob) as our chosen method of normalisation but reasoned that with the exception of Quantile and Funnorm, other normalisation methods would be applicable to MskAge.

### Existing Epigenetic Clocks are Inaccurate in Musculoskeletal Tissues Relative to a Bootstrapped Randomly Sampled CpG Model

We assessed the feasibility of using existing epigenetic clocks as age biomarkers in cartilage, bone, mesenchymal stromal cells (MSCs), skeletal muscle and tendon tissue. To draw a relative comparison, we compared errors of existing clocks to a “randomly sampled” model, which ran 2,500 elastic net regression models each on 350 randomly sampled CpGs from the training dataset and evaluated the predictions they made on independent test data. The distribution of the errors can be seen in Figure 2A, with a median absolute error of +/- 9.83 years for all tissues indicated by the dashed line. We compared the randomly sampled model errors and errors from existing epigenetic clocks averaged across all musculoskeletal tissues (Figure 2B) and split by each musculoskeletal tissue (Figure 2C). No existing epigenetic clock performs significantly better in terms of error than the randomly sampled model (Table 1). The SkinBlood clock had comparable performance with no statistically significant difference between errors (p = 0.31), all other epigenetic clocks had significantly greater errors than the randomly sampled model (p *<* 0.05) (Table 1).

**Fig. 2.**
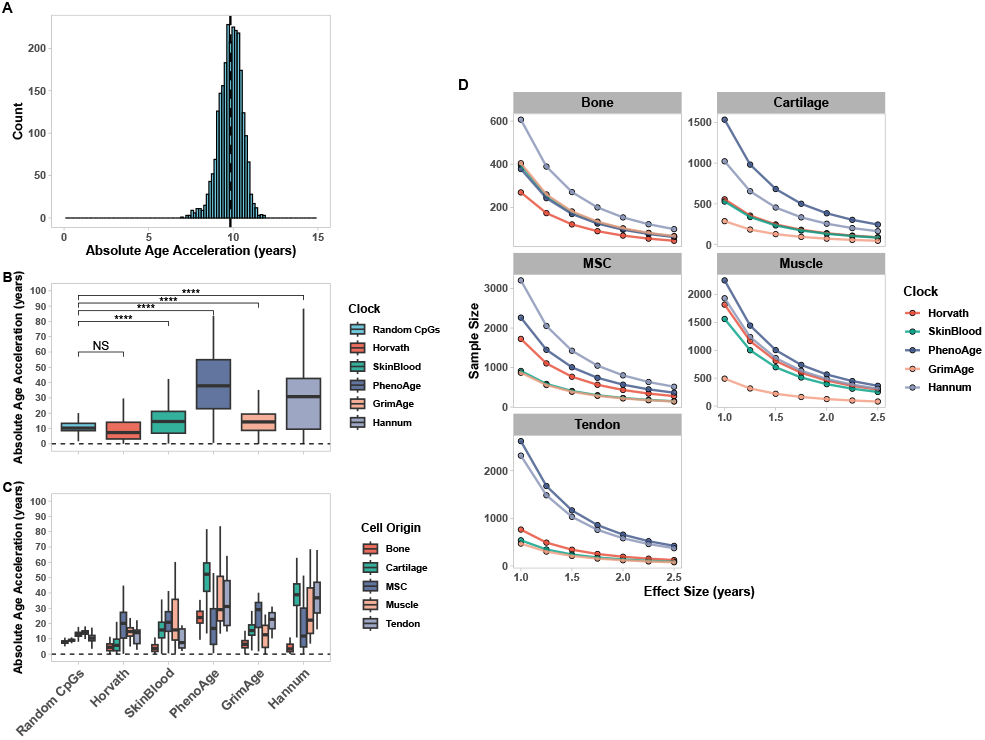
A random CpG model outperforms existing epigenetic clocks in musculoskeletal tissues. (A) Histogram displaying the distribution of absolute errors from 2,500 iterations of an elastic net regression model containing 350 randomly sampled CpGs evaluated on an independent test dataset, the dashed vertical line represents the median error (+/- 9.83 years). (B) Boxplots of absolute age acceleration derived from existing epigenetic clocks and the errors from the randomly sampled models. Boxplots display the combined errors across all musculoskeletal tissues (B) and the specific errors for each musculoskeletal tissue (C). Boxplots display median errors and the interquartile range. Units are expressed in years. (D) Power calculations derived from respective errors of each existing epigenetic clock for a range of given effect sizes 1 year to 2.5 years, the y axis refers to the sample size required to achieve 80% power for each respective effect size.

**Table 1.**
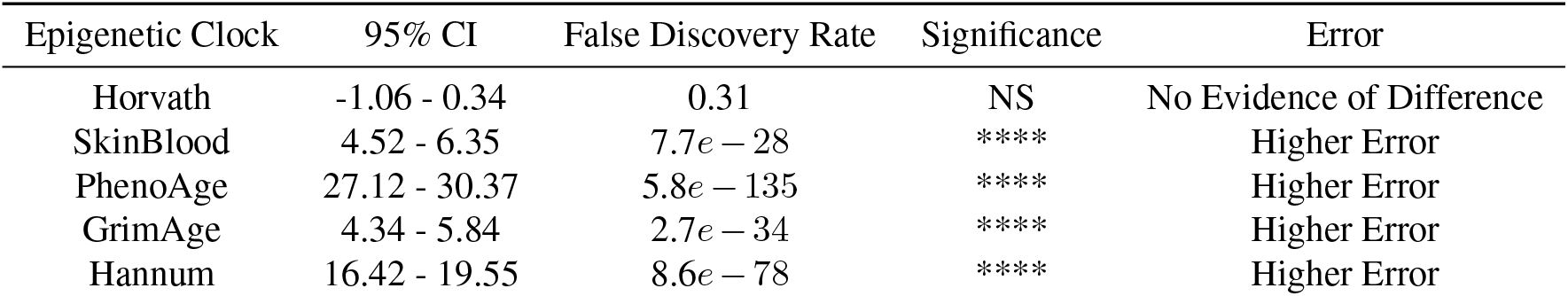
Confidence Intervals (CI) and False Discovery Rate (FDR) from pairwise Students T-tests comparing errors from existing epigenetic clocks to a model containing randomly sampled CpGs.

| Epigenetic Clock | 95% CI | False Discovery Rate | Significance | Error |
| --- | --- | --- | --- | --- |
| Horvath | -1.06 - 0.34 | 0.31 | NS | No Evidence of Difference |
| SkinBlood | 4.52 - 6.35 | $7.7e-28$ | **** | Higher Error |
| PhenoAge | 27.12 - 30.37 | $5.8e-135$ | **** | Higher Error |
| GrimAge | 4.34 - 5.84 | $2.7e-34$ | **** | Higher Error |
| Hannum | 16.42 - 19.55 | $8.6e-78$ | **** | Higher Error |

### Errors Associated with Existing Epigenetic Clocks Reduce the Capacity to Detect Perturbations in Epigenetic Ageing in Musculoskeletal Tissues

To further quantify the impact of errors from existing epigenetic clocks, we simulated power calculations for theoretical experiments. Power calculations were simulated based on detecting an effect size difference in that experiment of 1 year to 2.5 years at 80% power and a 5% false positive rate. Using existing epigenetic clocks, we demonstrate that the consequences of high errors diminsh the feasibility of designing appropriately powered *in vitro* experiments across most musculoskeletal tissues tested (Figure 2D). One example is that of MSCs, a commonly used cell type in tissue engineering that has attracted many efforts that attempt to characterise and modulate aspects of biological age (24, 25). Reliably detecting an effect size of 2.5 years in MSCs would require *>*200 samples using any of the existing epigenetic clocks.

### Development of a Highly Accurate Principal Component-Based Clock in Musculoskeletal Tissues

To build MskAge, we collected data from the public domain and within CIMA of 1,048 Infinium Methylation 450k or Infinium Methylation EPIC array samples that comprised multiple musculoskeletal tissues and blood. To eliminate bias, model development was performed on a training set (70%) and evaluation of error on an independent test set (30%). A penalised genetic algorithm islands model was designed to perform feature selection. MskAge was the result of regressing the principal components on the CpGs selected by the genetic algorithm against a transformed version of chronological age. The median absolute error across all tissues in the independent test data was +/- 3.51 years. The R^2^ of MskAge regressed against chronological age in the test set was 0.92 (Figure 3A & 3B). With the exception of tendon, which had a median absolute error of 5.52 years, all other tissues displayed a median absolute error of less than 4 years (Figure 3B). Moreover, we demonstrate that MskAge errors are consistent across tissue types when expressed as a percentage of chronological age (Supplementary Figure 3). The errors for each tissue are displayed in Table 2.

**Table 2.** Median absolute errors of MskAge test set predictions for each respective tissue and cell type in the test dataset

| Tissue | Median Absolute Error (years) |
| --- | --- |
| Blood | 2.91 |
| Bone | 3.32 |
| Cartilage | 3.92 |
| MSC | 3.25 |
| Muscle | 3.99 |
| Tendon | 5.62 |

**Fig. 3.**
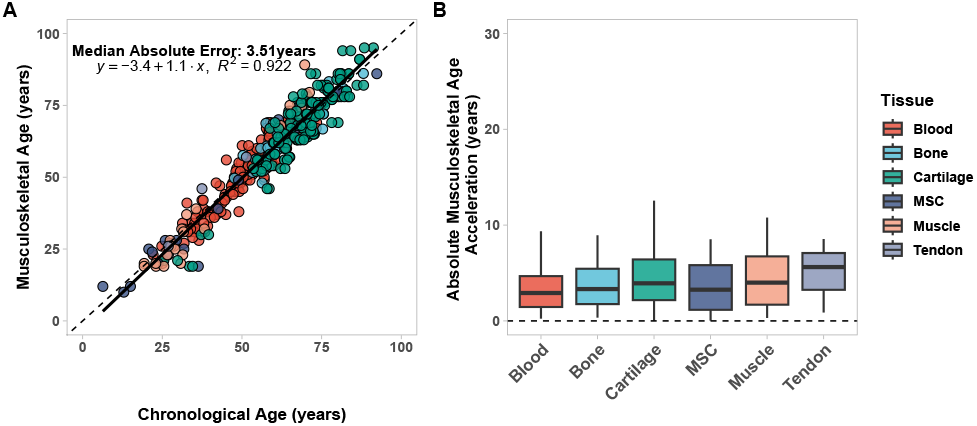
MskAge is a highly accurate predictor of age in musculoskeletal tissues and blood. (A) Linear regression of predicted epigenetic age generated with MskAge (y axis) vs chronological age (x axis) on independent test data (n=307). Points are coloured by tissue origin. (B) Boxplot of Median Absolute Errors (MAE) of epigenetic age acceleration and chronological age from predictions generated with MskAge on independent test data (n=307) split by tissue origin. Boxplots display median errors and the interquartile range. Units are expressed in years.

### CpGs used to construct MskAge are significantly enriched for terms related to skeletal and mesenchyme development

To further understand the biological context of methylation changes being captured by MskAge, we performed enrichment analysis for the genes that the CpGs in the model were located within. Using a network-based approach which connects nodes (significant enrichment terms, FDR *<* 0.05) via edges (gene-set similarity), we identify 4 clusters of Gene Ontology based enrichment terms (Figure 4). Notably, the largest cluster contained terms specifically related to skeletal, mesenchyme and connective tissue development. We employed the same approach using the Kyoto Encylopedia of Genes and Genomes (KEGG). Significant KEGG terms (FDR *<* 0.05) included those in cAMP, Hippo, Wnt and calcium signalling pathways and pluripotency of stem cells (Supplementary Figure 4).

**Fig. 4.**
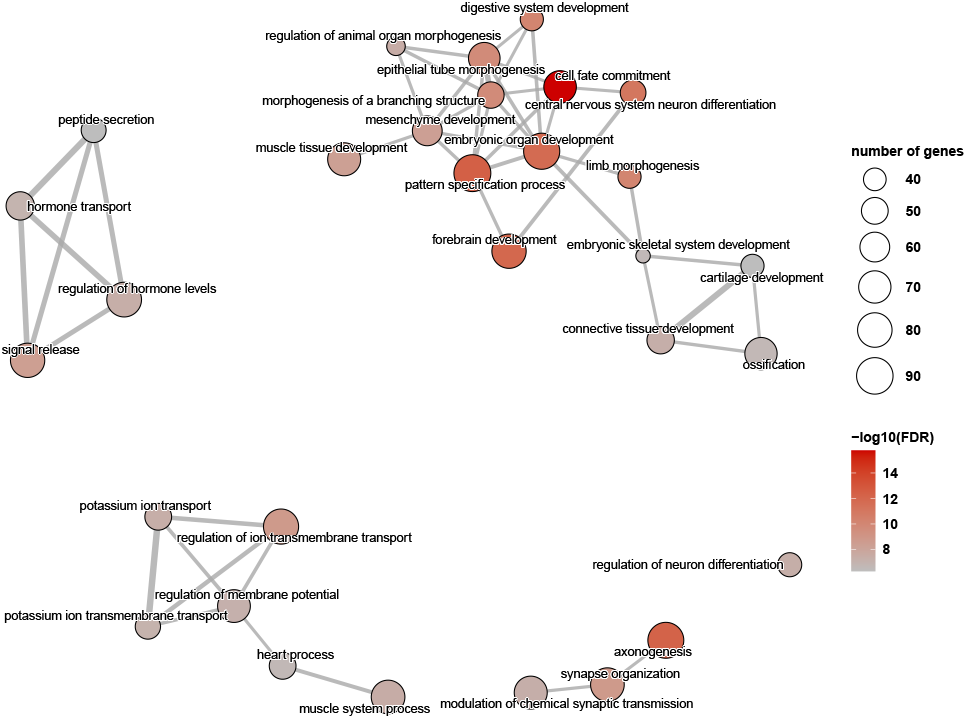
CpGs in MskAge are highly enriched for skeletal system and mesenchyme development. Enrichment map network of significantly enriched gene ontology terms for the individual coefficients of MskAge. The size of the nodes corresponds to the number of genes enriched within each gene ontology term. The colour of the node represents the -log10(FDR) of the enrichment for each term.

### The Musculoskeletal Clock is Reset with Cellular Reprogramming of MSCs and Fibroblasts

Given that our outcome variable was a transformed version of chronological age, a question that remains is whether or not MskAge is a highly accurate predictor of chronological age, or whether the biomarker has the capacity to track biological age. We utilised three *in vitro* models of cellular reprogramming to address this (26–28). We demonstrate that reprogramming of MSCs to iPSC-MSCs resets MskAge to approximately 0, consistent with the age observed in Embryonic Stem Cells (ESCs) (Figure 5A). Moreover, we observe the same age reduction over a longitudinal time course of fibroblasts being reprogrammed to iPSCs (Figure 5B). Importantly, MskAge was not trained on fibroblasts, but still has capacity to detect their rejuvenation as they are programmed towards iPSCs. Likewise, for ESCs, MskAge has not observed the specific methylation profile of an ESC but detects methylation in ESCs to have an epigenetic age of 0. Finally, we computed MskAge predictions on fibroblasts that were not fully reprogrammed to iPSCs but instead only reprogrammed until the maturation phase, a point at which the cell still retains its cellular identity and can redifferentiate (27) (Figure 5C). We observe that when fibroblasts are only partially reprogrammed, MskAge does not reset to 0 but instead decreases by approximately 30 years, which is in agreement with effect sizes observed by the Skin and Blood clock (27). It is widely accepted that cellular reprogramming is one of the most potent methods of cellular rejuvenation, thus highlighting the capacity for MskAge to track underlying changes in biological ageing.

**Fig. 5.**
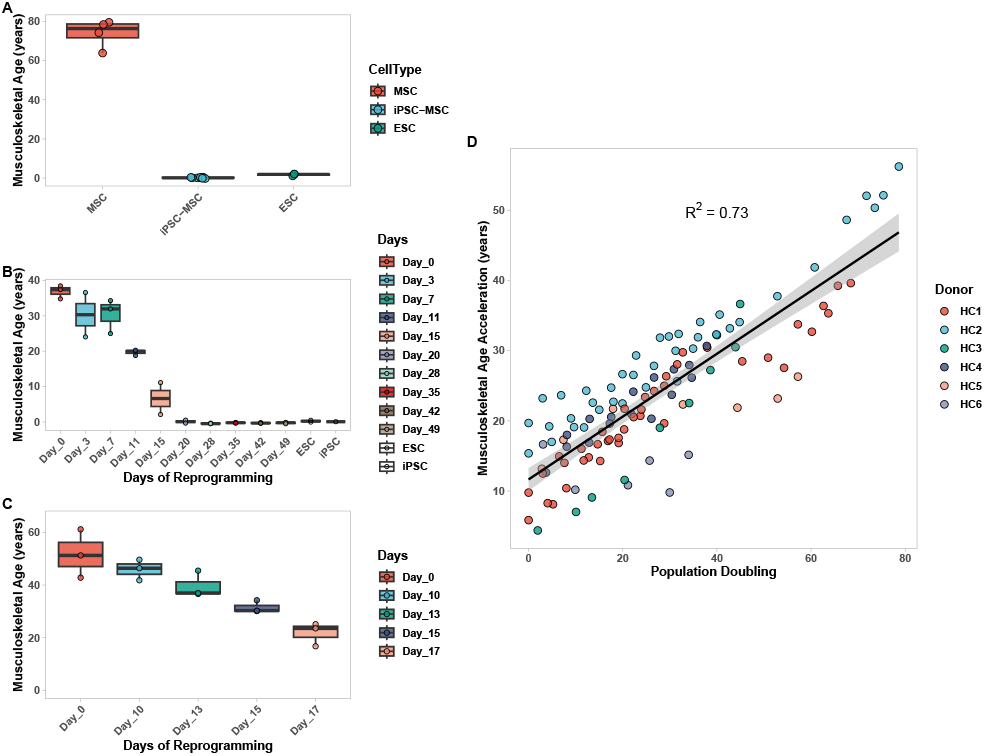
MskAge tracks cellular ageing and rejuvenation *in vitro*. (A) Linear regression of predicted epigenetic age generated with MskAge (y axis) vs population doubling of fibroblasts from 6 healthy donors (x axis). (B) Boxplot of predicted epigenetic ages generated with MskAge of MSC donors and the same cells transformed to iPSC-MSCs. Epigenetic age predictions were also computed on ESCs from the same dataset as a negative control. (C) Boxplots of predicted epigenetic ages generated with MskAge on human dermal fibroblasts over a longitudinal experiment of reprogramming to iPSCs. Epigenetic age predictions were also computed on ESCs from the same dataset as a negative control. (D) Boxplots of predicted epigenetic ages generated with MskAge on human dermal fibroblasts following a longitudinal experiment of iPSC reprogramming only to the maturation phase.

### Musculoskeletal Age tracks *in vitro* cellular ageing

We envision that one of the prominent use cases for MskAge will be to facilitate in vitro research on musculoskeletal cells. Thus, a key attribute of such a tool would be that it tracks agerelated methylated changes *in vitro* as well as it does on *ex vivo* tissues. We sought to test whether MskAge tracks epigenetic ageing in vitro over multiple cell divisions across a longitudinal in vitro time course. We reprocessed and analysed the Cellular Lifespan multi-omics longitudinal fibroblast ageing study (29) that includes six healthy control fibroblast cell donors cultured for up to 80 population doublings. We calculated MskAge and extracted epigenetic age predictions of existing clocks on the same data. MskAge acceleration exhibits the strongest linear relationship (R2 = 0.73, FDR *<* 2.37e-34) with population doublings relative to all clocks tested, including the fibroblast-specific SkinBlood clock (R2 = 0.51, FDR *<* 1.99e-15) (Figure 5D). According to MskAge in this study, *in vitro* ageing occurs at an average rate of 0.45 years per population doubling (Table 3). The ability of MskAge to detect *in vitro* age acceleration as a product of population doubling is advantageous for its use in facilitating the identification of perturbations that can attenuate or accelerate the ageing process in musculoskeletal cells.

**Table 3.** Output of linear regression models computed from epigenetic age predictions in of MskAge and epigenetic clocks in the Longitudinal Cellular Lifespan Study. PC stands for the Principal Component version of the clocks.

| Clock | R<sup>2</sup> | P Value | FDR | Intercept | Slope |
| --- | --- | --- | --- | --- | --- |
| **MskAge** | **0.73** | **2.15e-35** | **2.37e-34** | **11.64** | **0.45** |
| Horvath | 0.05 | 0.031 | 0.124 | 18.84 | 0.17 |
| SkinBlood | 0.51 | $1.99e-15$ | $1.79e-14$ | -7.27 | 0.33 |
| PhenoAge | 0.05 | 0.033 | 0.124 | 3.09 | 0.20 |
| GrimAge | 0.03 | 0.079 | 0.158 | 26.82 | -0.04 |
| Hannum | 0.66 | $6.90e-23$ | $6.9e-22$ | -36.40 | 0.41 |
| PCHorvath | 0.18 | $2.58e-05$ | $2.0e-4$ | 23.08 | 0.14 |
| PCSkinBlood | 0.11 | 0.001 | 0.005 | 11.37 | 0.16 |
| PCPhenoAge | 0.40 | $1.25e-11$ | $1.0e-10$ | 59.21 | 0.27 |
| PCGrimAge | 0.02 | 0.165 | 0.166 | 52.75 | 0.03 |
| PCHannum | 0.39 | $3.77e-11$ | $2.64e-10$ | 36.77 | 0.22 |

### Musculoskeletal Age is Accelerated in Lesioned Osteoarthiritic Cartilage

MskAge was trained on nonlesioned cartilage samples. To test whether MskAge is accelerated in the context of Osteoarthiritis (OA) we reprocessed 298 hip and knee non-OA and OA cartilage samples. Lesioned cartilage exhibits significant Musculoskeletal Age acceleration in the hip (4.029 years, FDR = 0.003) and borderline significant Musculoskeletal Age acceleration in the knee (3.16 years, FDR = 0.067) relative to preserved non-OA controls (Figure 6 A & B) (Table 4). Interestingly, cartilage extracted from non-lesioned hip OA patients still exhibits a trend towards being epigenetically older than preserved control hip cartilage (2.056 years, p = 0.094) (Figure 2A) (Table 4). While the direction of change is also the same for preserved OA knee cartilage relative to preserved control knee cartilage (1.02 years) the difference is non-significant (p = 0.231). It is noteworthy that no significant differences in MskAge were observed when we compared healthy to osteopenic and osteoporotic bone samples (Supplementary Figure 5).

**Table 4.** Output of pairwise students t-tests for comparisons of epigenetic age predictions derived from MskAge in hip and knee cartilage samples. A positive mean difference indicates epigenetic age acceleration in the subgroup listed on the right hand side of the contrast

| Contrast | P Value | FDR | Mean Difference (years) |
| --- | --- | --- | --- |
| Preserved Control Hip vs Lesioned OA Hip | $4.7e-4$ | 0.003 | 4.029 |
| Preserved OA Hip vs Lesioned OA Hip | 0.104 | 0.128 | 2.056 |
| Peserved Control Hip vs Preserved OA Hip | 0.047 | 0.094 | 1.974 |
| Preserved Control Knee vs Lesioned OA Knee | 0.022 | 0.067 | 3.16 |
| Preserved OA Knee vs Lesioned OA Knee | 0.107 | 0.128 | 2.14 |
| Peserved Control Knee vs Preserved OA Knee | 0.231 | 0.231 | 1.02 |

**Fig. 6.**
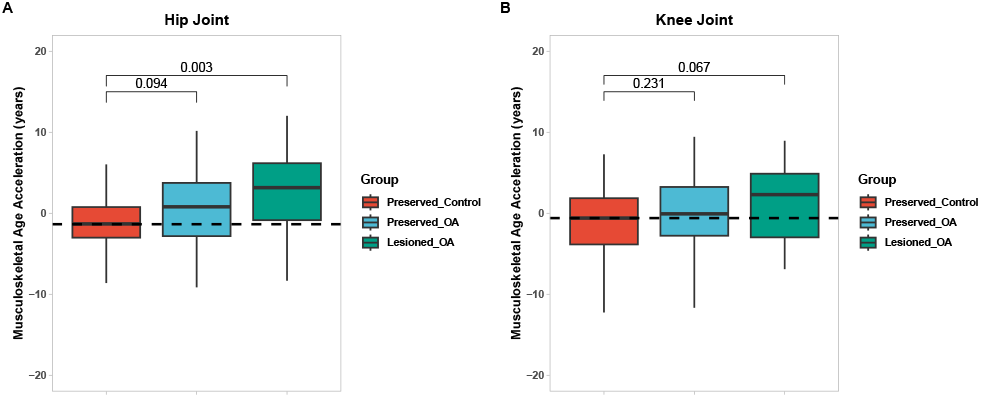
MskAge is accelerated in lesioned OA cartilage. (A) Predicted MskAge of Preserved Control (n=73), Preserved OA (n=39) and Lesioned OA (n=26) Hip cartilage samples. (B) Predicted MskAge of Preserved Control (n=58), Preserved OA (n=85) and Lesioned OA (n=17) Knee cartilage samples.

## Discussion

The musculoskeletal system is crucial to the ageing process, and maintaining its proper function is key for extending both health span and lifespan in humans (30). Although there has been extensive research on the ageing of the musculoskeletal system, few reliable biomarkers are available for musculoskeletal tissues (18). In this study, we introduce MskAge, an accurate principal component-based epigenetic clock that can be applied *ex vivo* and *in vitro* to investigate the epigenetic age of musculoskeletal tissues and cells. MskAge tracks biological age acceleration and reversal, making it a promising tool for studying pro or anti-ageing interventions in musculoskeletal tissues and cells.

Recent work has emphasised the importance of preprocessing pipelines for epigenetic age prediction (31). In light of this, we evaluated important technical considerations during the construction of MskAge by assessing the impact of various array-based normalisation methods on epigenetic age predictions. A previous study demonstrated that Horvath’s age prediction is robust to the choice of data preprocessing methods (32). While our results align with McEwen and colleagues, the same does not hold true for other clocks, particularly when employing quantile-based normalisation methods for PhenoAge, GrimAge and Hannum. Quantile normalisation methods force data distributions to be equal, resulting in a stringent method of statistical normalisation. Notably, PhenoAge, GrimAge and Hannum clocks did not have quantile normalised data in their training sets, which may account for the exacerbated prediction errors. As a result, we recommend normalising data using ssNoob prior to making predictions with MskAge.

A common debate in the ageing biomarker literature concerns the absolute importance of accuracy when the outcome variable is chronological age (33–35). It is indeed valid that outside of forensic settings, establishing a model that provides a perfect readout of chronological age is seldom useful. Zhang and colleagues show experimentally that as a blood-based epigenetic clock’s predictions approach near-perfect chronological age accuracy, the clock loses its association with mortality risk, which is a proxy of biological age (35). These findings create some subjectivity and uncertainty around the ideal level of accuracy for a biomarker to be useful.

To address this question objectively, we used a two-fold datadriven approach to assess the applicability of existing epigenetic clocks were applicable to musculoskeletal tissues. First, we created a “Random CpG” model, involving 2,500 iterations of randomly selecting 350 CpGs from the 338,185 CpGs in the dataset, training a model on 70% of the data and evaluating its error on the 30% held-out test set. We selected 350 CpGs because it is approximately equal to the mean number of CpGs in the existing epigenetic clocks tested. Remarkably, the errors from the “Random CpG” model were as accurate or, in most cases, more accurate than epigenetic age predictions using existing clocks on the same samples (Figure 2B,2C, Table 1). This outcome underlines the extent to which existing epigenetic clock CpGs lack specificity for musculoskeletal tissues. A plausible explanation for the accuracy of the Random CpG model is that it reflects the degree to which ageing impacts methylation patterns across a substantial portion of the epigenome, which has been reported to be up to 20% (36). Accurate models of chronological age prediction have also been built using only a small number of CpGs, such as those in the genes ELOVL2, FHL2, KLF14 and TRIM59 (37).

Secondly, we simulated power calculations based on the errors of existing epigenetic clocks across a range of theoretical effect sizes for future experiments. We propose that one of the primary applications for an epigenetic biomarker of age in musculoskeletal tissues would be to identify compounds that could perturb the ageing process *in vitro*. According to the power calculations, designing adequately powered studies using existing epigenetic biomarkers would be largely impractical and unfeasible both logistically and financially. The combined evaluation of both above methods objectively underscores that existing epigenetic clocks are not suitable for quantifying epigenetic ageing in musculoskeletal tissues. As a result, we developed MskAge with the goal of providing substantial improvements in predictive accuracy and musculoskeletal-tissue specificity compared to other epigenetic clocks. Furthermore, we benchmarked MskAge to show that its enhanced age prediction accuracy did not compromise its ability to detect changes in biological age.

Developing a multi-tissue epigenetic clock can be viewed as a multi-objective optimisation problem. DNA methylation serves as a critical factor in determining cellular identity, and as a result, differences in methylation between tissues tend to be more pronounced than the more subtle changes occurring with age (38). The creation of a multi-tissue epigenetic clock adds a hierarchical structure to a dataset. When age distributions across various tissue types are unbalanced, a naïve model that overlooks tissue origin might inadvertently capture tissue-specific methylation patterns rather than age-related changes.

MskAge was developed to address this issue by employing a multi-level genetic algorithm islands model that minimises within-tissue age prediction errors. Genetic algorithms are meta-heuristic frameworks that optimise outcomes based on evolutionary principles (39). We utilised a penalised ridge regression fitness function within the genetic algorithm framework to efficienctly search and reduce the feature space. Ridge regression models have been successfully employed as learners in fitness functions for genetic algorithms across various applications (40, 41). The L2 penalty of the ridge regression cost function causes the coefficient weights of correlated features to shrink towards each other without completely removing them (42). This coefficient shrinkage is desirable in the context of CpG methylation, which exhibits high collinearity. Furthermore, since the coefficient weights are shrunken but not removed, as in lasso or elastic net, ridge regression ensures that the inclusion of CpGs is assigned at some weight, enabling them to be penalised for their inclusion in the model.

To penalise the inclusion of CpGs, we assigned each CpG a penalty of 0.01 years in the genetic algorithm, promoting a fitness benefit that encourages convergence towards a model with fewer features, provided that the removal of a specific CpG does not inflate the model error by more than the specified penalty. Rather than a single genetic algorithm, we distributed the evolutionary optimisation task across four independent islands that exchanged information on their best solutions every ten iterations. Utilising multiple islands for the optimisation task enables a more efficient search of the feature space and diminishes the likelihood of converging on local optima (43, 44).

After 200 iterations, our “fittest” model, which includes the selection of CpGs optimised by the genetic algorithm to achieve the lowest cross-validated accuracy, contained 3,365 CpGs. Rather than further reducing this selection, we transform all 3,365 CpGs into linear principal components using Singular Value Decomposition (SVD). The use of using principal components as a dimensionality reduction method in constructing multivariate models with ‘omics data is not new (45, 46). However, the first attempt using this approach in the development of epigenetic clocks was published recently (19). What distinguishes our method from Higgins-Chen and colleagues is we adapted this approach by performing SVD on M values after extensive variable selection through the genetic algorithm. This approach resulted in an improved test set error of +/- 3.51 years, compared to an error of +/- 5.21 years when creating a principal component model on the full CpG matrix before variable selection.

The advantages of using principal components as features in epigenetic clocks have been discussed previously (19). An epigenetic clock constructed on principal components of a larger number of age-related CpGs improves both inter- and intra-dataset predictions, addressing the significant challenge of array-related “batch to batch” variability (19). Taking such an approach means that MskAge is optimised for data generated from Infinium 450 or Infinium EPIC arrays due to the large amount of CpGs that need to be quantified. Although it is feasible to develop a subset of MskAge coefficients that explain maximum variance with minimal redundancy, this likely would reduce the overall accuracy of the model.

To date, little mechanistic evidence has emerged from epigenetic clock coefficients, and it has often been challenging to infer functional relevance to the ageing process from CpG-based models containing a few hundred CpGs (33). The highly co-linear nature of the epigenome may inflate the chances of clocks being built on methylation changes that are correlated but not causal to the ageing process. The method employed in this study does not eliminate the inclusion of correlated, non-causal features. However, we found that by including a larger number of correlated CpGs, there was significant enrichment for Gene Ontology processes associated with muscle, cartilage, connective tissue, mesenchyme, and skeletal system development. Moreover, the enriched KEGG pathways identify key signalling pathways, such as Hippo, Calcium and Wnt, which are commonly implicated in both musculoskeletal ageing and disease. Wnt signalling regulates numerous cellular functions and importantly contributes to both bone and cartilage regeneration, and its dysregulation exacerbates osteoarthritis (47). Additionally, hippo signalling has been implicated in regulating skeletal muscle mass (48). Interestingly, both Gene Ontology and KEGG terms contain modules related to hormonal transport and Cushing syndrome, respectively. Cushing syndrome is a condition related to the overproduction of cortisol, and hypercortisolism is proposed to induce a premature ageing phenotype (49). While further research is required to delineate the functional relevance of methylation changes in these pathways, the enrichment of MskAge coefficients in musculoskeletal and ageing-specific pathways is promising.

We assessed MskAge’s capacity to quantify biological ageing methylation changes using cellular reprogramming and longitudinal *in vitro* ageing. Cellular reprogramming, the gold standard of experimental cellular rejuvenation, reverts somatic cells into an ESC-like state, reversing many ageing hallmarks and rejuvenating the transcriptome and methylome (7, 26–28, 50). We reanalysed three independent cellular reprogramming datasets, demonstrating that MskAge consistently reports the age of iPSCs and ESCs as zero and tracks the biological reversal of fibroblasts and MSCs during reprogramming, even though these some of these cell types were not included in MskAge’s training (26–28).

It is plausible that by using the principal components of a large number of CpGs (3,365), MskAge has the scope to capture cell-type specific and cell-type independent features of biological ageing. In contrast to cellular reprogramming, serially passaged primary cells without genetic perturbations undergo systemic changes recapitulating ageing and eventually become senescent (51). MskAge exhibited a strong linear relationship with population doubling in serially passaged fibroblasts, outperforming other clocks tested, including the SkinBlood clock (4). MskAge’s ability to track *in vitro* ageing is advantageous for musculoskeletal ageing research, which often relies on the extraction and culture of *ex vivo* cells. Furthermore, MskAge ticks at an approximate rate of 0.45 years per population doubling in primary fibroblasts, enabling researchers to objectively investigate perturbations that may modulate this rate.

MskAge constitutes a valuable new tool for musculoskeletal biologists and ageing researchers that is a highly accurate predictor of epigenetic age in musculoskeletal cells and tissues *ex vivo* and *in vitro*. Its superior accuracy facilitates the design of experiments with appropriate statistical power compared to existing epigenetic clocks in musculoskeletal tissues. In addition to its precision, MskAge effectively tracks the biological age of cells through well-known ageing perturbations and provides a means by which serial passaging of cells can be used as a model of epigenetic ageing *in vitro*. MskAge also facilitates the monitoring of pharmacological, genetic, and nutritional interventions on musculoskeletal cells.

## Methods

### Data Acquisition

Datasets used for the development of the Musculoskeletal Clock were acquired internally from the Center for Integrated Musculoskeletal Ageing (CIMA) or the public repository Gene Expression Omnibus (GEO). GEO search terms were restricted to data generated on the Infinium Methylation 450k or Infinium Methylation EPIC array platforms. Sources and descriptions of the acquired data can be found in Supplementary Table 1.

### Data Processing

All data processing was performed in R (4.1.2). Raw IDAT files and raw methylated and unmethylated intensity matrices available for 16/18 datasets were reprocessed using the functionality within the minfi package (20). CpG probes were removed from IDAT files if their methylated and unmethylated signal intensities were not significantly above that of control probes (detection p-value *>* 0.01). Probes were further removed if they mapped within 2 base pairs of a single nucleotide polymorphism (SNP), to the X or Y chromosome, or were found to cross-hybridise to multiple genomic locations (52, 53). The remaining 2/18 datasets were processed as above with the exception that signal intensity relative to control probes could not be assessed.

### Epigenetic Age Calculations for Existing Epigenetic Clocks

All epigenetic age calculations for existing epigenetic clocks presented in this manuscript were calculated using the online DNA Methylation Age calculator with the “Normalize Data” option selected (http://dnamage.genetics.ucla.edu/). Age calculations for the in vitro fibroblast ageing data were derived directly from the Cellular Lifespan Study shiny application Cellular Lifespan Study (29). It is valuable to note that whilst two iterations of a skeletal muscle clock have been published, they could not be evaluated herein because all of the skeletal muscle samples used in this study were used in the training sets of the two iterations of the skeletal muscle clock (8, 54).

### Data Normalisation

Following the investigation of various normalisation methods, datasets used in the development of the musculoskeletal clock for which raw IDAT files were available were normalised using single sample normal-exponential out-of-band (ssNoob) normalisation (21). Datasets for which only unnormalised beta values were available were normalised using Beta Mixture Quantile Normalisation (BMIQ) (23).

### Data Imputation

Probe-wise and sample-wise missingness were assessed by counting row-wise and column-wise missing values (NA’s). The data was overall complete, with *>*99.5% of samples having missing values for *<*1% of their CpGs. Values for the CpGs that were missing were imputed using the impute.knn function (55).

### Defining a random CpG model of DNAmAge in Musculoskeletal Tissues

To establish a random CpG model, the dataset was split into training (70%) and test data (30%) with respect to tissue origin for the samples. A 10-fold cross-validated elastic net regression model was trained using 350 CpGs sampled at random without replacement from the 338,185 CpGs that remained after filtering and integrating the data. This process was repeated 2,500 times to provide an unbiased distribution of errors (56). The error was calculated as the absolute Age Acceleration (DNAmAge – Age).

### Determining Power Calculations for Given Effect Sizes

For each respective error of existing clocks in musculoskeletal tissues, power calculations were simulated based on a fixed power of 80% and significance at 5% (p=0.05). Cohen’s d was calculated as the difference in means divided by the pooled standard deviation of DNAm Age Acceleration. Power calculations were simulated for effect sizes of 1 year to 2.5 years in increments of 0.25 years.

### Univariate statistical analysis

Univariate statistical tests were computed with either pairwise student’s t-tests or linear models as described in the table and figure legends. Resulting p values were adjusted using the Benjamini Hochberg FDR method (57).

### Genetic Algorithm

#### Training and Test Set Splits

Data was split 70/30 into training and test sets with respect to tissue origin. Descriptions of samples within each set can be seen in Table 5.

**Table 5.** Training and Testing for development of MskAge.

| Set | Female | Male | Total | Age Range (years) |
| --- | --- | --- | --- | --- |
| Training | 468 | 270 | 738 | 2 - 95 |
| Test | 169 | 136 | 305 | 10 - 95 |

**Table 6.** Genetic Algorithm Parameters.

| Parameter | Value |
| --- | --- |
| Iterations | 200 |
| Population Size | 200 |
| Islands | 4 |
| Crossover | 0.7 |
| Crossover Method | Uniform Crossover |
| Mutation Rate | 0.1 |
| Mutation Method | Uniform Random Mutation |
| Migration | 10 |
| Migration Rate | 0.1 |
| Elitism | 2 |

**Table 7.** Data sources used to build MskAge

| Tissue/Cell Type | Sample No | Source |
| --- | --- | --- |
| Blood | 544 | GSE128235, GSE42861 |
| Bone | 90 | GSE202067, PMID: 29514638 |
| Cartilage | 255 | GSE63695, CIMA Dataset(s) |
| Mesenchymal Stromal Cells | 69 | GSE202067, GSE52114, GSE79695, CIMA Dataset |
| Skeletal Muscle | 82 | GSE171140, GSE151407, GSE50498 |
| Tendon | 8 | CIMA Dataset |

**Table 8.** Patient/Sample characteristics used to build MskAge

| CellType | Age | Males | Females |
| --- | --- | --- | --- |
| Blood | 51 ± 12.4 | 180 | 364 |
| Bone | 64.4 ± 9.7 | 2 | 88 |
| Cartilage | 69.0 ± 13.9 | 106 | 149 |
| MSC | 39.1 ± 27.5 | 30 | 34 |
| Muscle | 40.5 ± 22.3 | 82 | 0 |
| Tendon | 40.8 ± 24 | 6 | 2 |

#### Feature reduction and selection

To reduce dimensionality by removing potentially redundant features (CpGs) we calculated the Pearson correlation coefficient of each CpG with age across each tissue in the training data and filtered out any CpGs with an absolute Pearson correlation *<*0.3. After filtering, 7,677 CpGs remained. We employed a penalised genetic algorithm islands model to select CpGs that were most predictive across each musculoskeletal tissue. A description of the algorithm can be found in sections below.

#### Problem Description

Using the reduced set of CpGs, the aim was to develop a model that optimised the selection of CpGs for the prediction of age across multiple musculoskeletal tissues and blood that also accounted for the hierarchical nature and imbalance between tissues in the training dataset. To do this, we designed a genetic algorithm islands model as follows.

Given a set of training samples Z *Z*_1_, *Z*_2_…*Z*_*i*_ with each *Z*_*i*_ having a corresponding chronological age *A*_*i*_, tissue original T *T*_1_, *T*_2_…*T*_*k*_ and an associated vector of M values for CpGs S *S*_1_, *S*_2_…*S*_*j*_. For convenience, we add a superscript index to each sample that denotes the tissue type that the sample belongs for. For example, *Z*_*t*_ denotes all samples in Z that belong to tissue T. The aim is to select a combination of S that minimises the error derived from the fitness function of predicting age *A*^*T*^ in samples *Z*^*T*^.

#### Encoding the fitness function

To select a subset of S predictive of age across all *Z*^*T*^, each CpG *S*_*i*_ is binary encoded to a value of *S*^1^ or *S*^0^. Here we denote a chromosome C as the binary encoded selection of S for a particular iteration of the genetic algorithm, specifically *S*^1^ ∈ *S*. To define the fitness (accuracy) of each chromosome C, we build a ridge regression function with lambda parameter defined by 10-fold cross-validation in the training set. The ridge regression fitness function can be seen below.

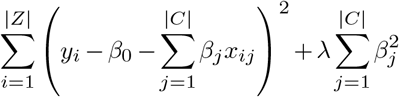

Using the ridge regression coefficients, 10-fold cross-validation on the training set Z is used to compute a mean absolute error (MAE) for each musculoskeletal tissue *Z*^*T*^. The MAE for each tissue is then averaged to calculate a single objective fitness value which is equal to the mean MAE equally weighted for every tissue T in the training set. We penalise the inclusion of CpGs in the model by adding a weight of 0.01 years to the MAE, thus forcing the genetic algorithm to discard individual CpGs whos inclusion in the model does not reduce error by at least this amount.

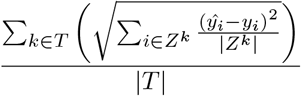

#### Evolution of chromosomes on independent islands

The task of minimising the fitness value by the selection of the fittest chromosomes is distributed across four different islands within the genetic algorithm. Initially, a population that contains 200 randomly sampled chromosomes is generated and split equally across eight island nodes. Each island evaluates its population of chromosomes as described above, resulting in 50 fitness values. The top 5% of chromosomes that yield the fittest individuals (i.e. those with the lowest fitness values) in each island are retained from that population. Single-point crossover is applied to occur at a rate of 70% to the top 5% of fitness individuals in each island. Crossover produces child chromosomes that are the product of two parent chromosomes. Random mutations are permitted to occur in each of the child chromosomes probabilistically at a rate of 10%. The aforementioned process is repeated for 400 iterations, with the migration of the fittest chromosomes between islands every ten iterations.

### Development of the Final Musculoskeletal Clock

To build the final model, we used the 3,365 CpGs selected by the genetic algorithm to compute principal components. We trained an elastic net regression model on these principal components to predict a transformed version of age (7). Age transformation was performed as follows.

If age *<*20:

F(age) = log(age+1)-log(20+1)

Else:

F(age) = (age-20)/(20+1)

To assess the accuracy of the model on hold-out test data and new independent datasets, M values were first projected into the same principal component space as the training data. The resulting principal components were then used to make predictions with the model.

### Functional Enrichment

Functional enrichment of the 3,365 CpGs was conducted using the clusterProfiler R package (58). CpGs were mapped to genes based on the Illumina manifest file. Enrichment was performed using the enrichGO function that performs enrichment of a vector of genes within Gene Ontology terms. Redundant terms were removed using the simplify function with a similarity cutoff of 0.7. Significant gene ontology pathway networks were clustered and visualised using an Enrichment Map. Edges connecting the nodes denote the overlap in genes between the pathways.

## ACKNOWLEDGEMENTS

We would like the acknowledge Dr Arturas Grauslys and the Computational Biology Facility at the University of Liverpool for expert statistical discussion during model development stages. Moreover, we would like to acknowledge Dr Mohammad Saniee Abadeh for advice conceptualising the genetic algorithm approach.

## Supplementary Note 1: Supplementary Methods

### Data Acquisition

Samples used to build MskAge were acquired from a combination of internal Center for Integrated Musculoskeletal Ageing (CIMA) datasets and a search of the public data repository Gene Expression Omnibus (GEO). The search terms for GEO were restricted to include Infinium 450 and Infinium EPIC arrays only. Search criteria allowed for the inclusion of any cells or tissues that function as part of the musculoskeletal system and blood. Given the abundance of publicly available blood methylation datasets, we decided to restrict the inclusion to no more than approximately half of the samples collected. Where feasibly possible, we used only tissue samples that were not diseased, although it was not plausible to obtain a cohort that was free of other co-morbidities. For cultured cells, such as Mesenchymal Stromal Cells (MSCs), we utilised only control cells from the experiments and filtered out samples that were exposed to any pharmacological, nutritional or genetic manipulation. All skeletal muscle samples were obtained from the *vastus lateralis*. Two out of the three skeletal muscle datasets (GSE171140, GSE151407) were repeated measures designs of individuals subjected to exercise programs. We used only baseline samples from these experiments. A summary of samples and the data sources can be seen in Supplementary Table 1.

**Fig. 7.**
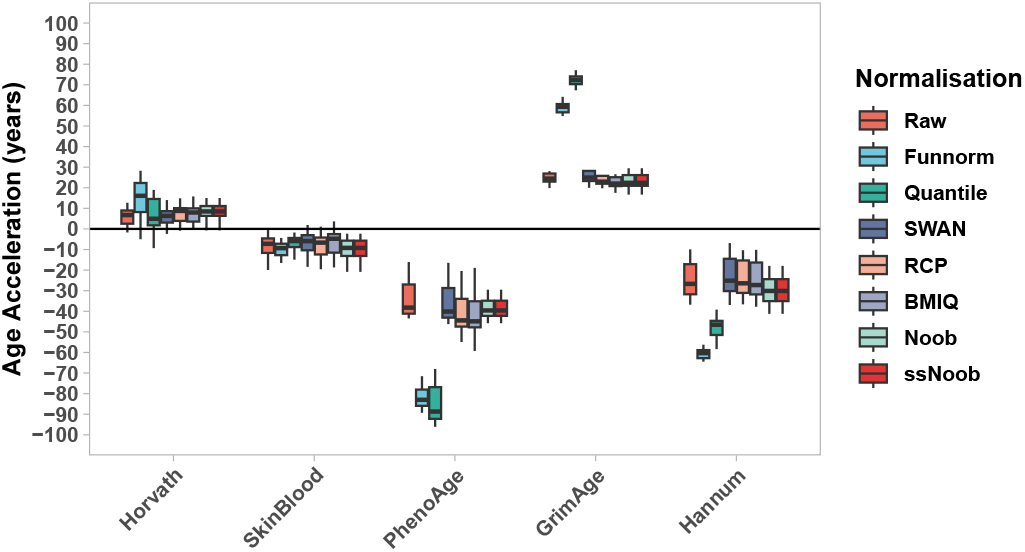
Supplementary Figure 1. Age acceleration of 16 cartilage samples (50-60 years) for Horvath, SkinBlood, PhenoAge, GrimAge and Hannum epigenetic clocks each normalised with raw (no normalisation), functional normalisation (Funnorm), quantile normalisation (Quantile), subset-within array normalisation (SWAN), regression of correlated probes normalisation (RCP), beta mixture quantile normalisation (BMIQ), normal exponential using out-of-band probes (Noob) and single-sample normal exponential using out-of-band probes (ssNoob). Units are displayed in years.

**Fig. 8.**
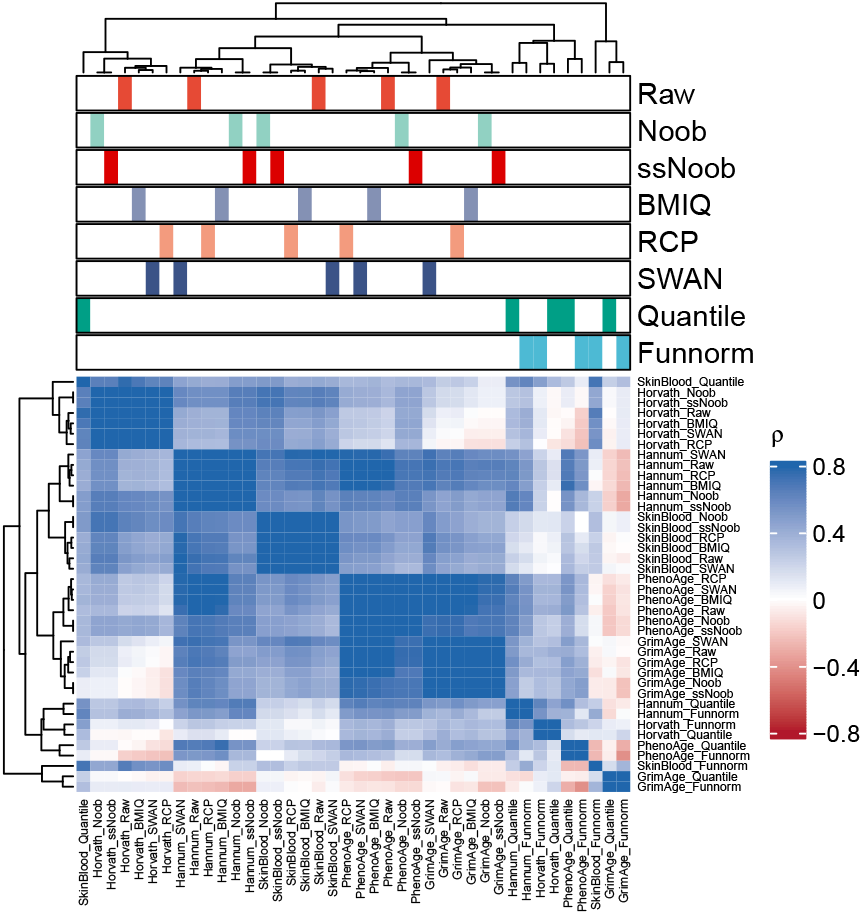
Supplementary Figure 2. Correlation heatmap of 16 cartilage samples (50-60 years) for Horvath, SkinBlood, PhenoAge, GrimAge and Hannum epigenetic clocks each normalised with raw (no normalisation), functional normalisation (Funnorm), quantile normalisation (Quantile), subset-within array normalisation (SWAN), regression of correlated probes normalisation (RCP), beta mixture quantile normalisation (BMIQ), normal exponential using out-of-band probes (Noob) and single-sample normal exponential using out-of-band probes (ssNoob). Correlation is spearman’s rho. Clustering was performed using Euclidean distance with the ward.D2 linkage method.

**Fig. 9.**
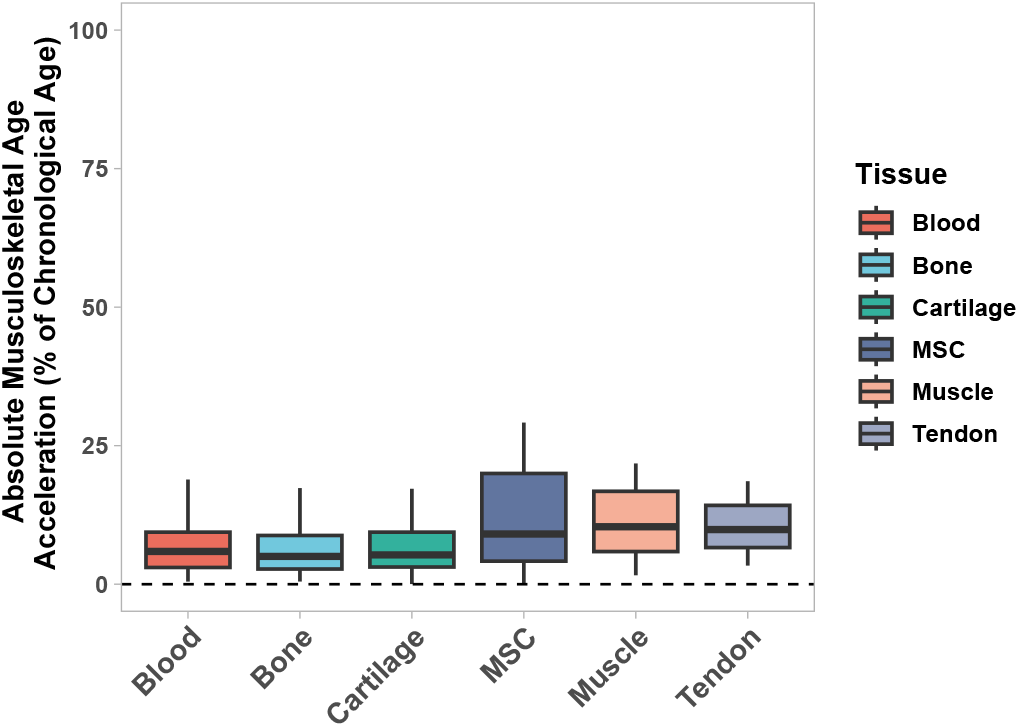
Supplementary Figure 3. Boxplot of Median Absolute Errors (MAE) from predictions generated with MskAge on independent test data (n=307) split by tissue origin and expressed as a percentage of chronological age.

**Fig. 10.**
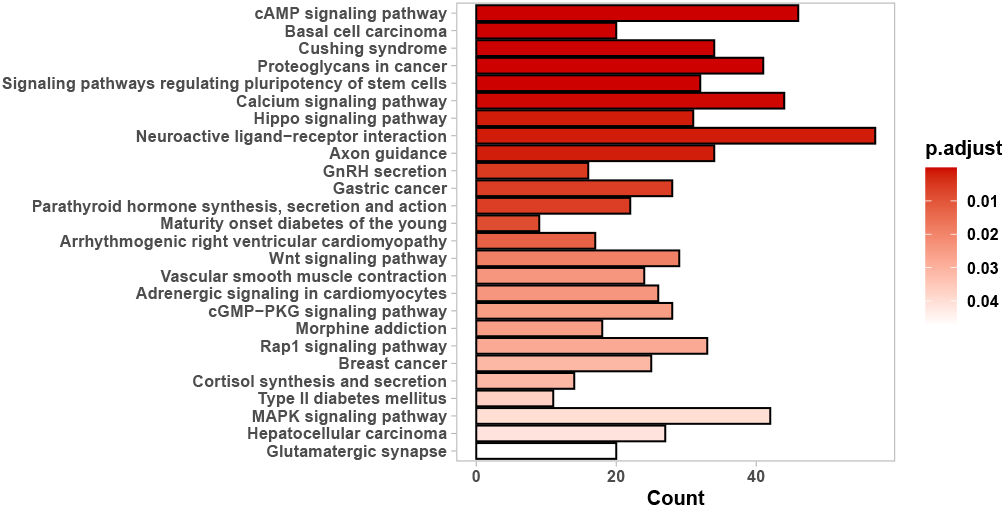
Supplementary Figure 4. Barplot of significantly enriched KEGG terms of genes proximal to MskAge. Bars are coloured according to their level of significance for each term.

**Fig. 11.**
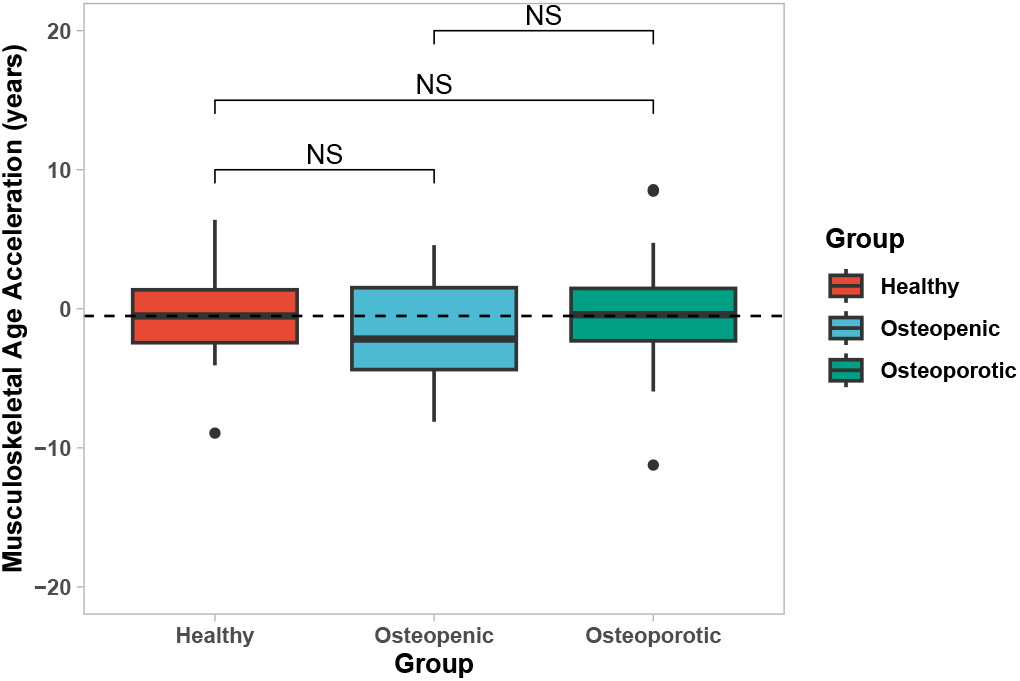
Supplementary Figure 4. Boxplot of Musculoskeletal Age Acceleration (years) from predictions generated with MskAge on bone biopsy samples from healthy (n=40), osteopenic (n=18) and osteoporotic (n=26) patients.

